# A unified framework for drug–target interaction prediction by semantic-guided meta-path method

**DOI:** 10.1101/2024.10.14.618129

**Authors:** Haoyuan Li, Jianxin Wang, Haochen Zhao, Kai Zheng, Qichang Zhao

## Abstract

Drug-target interaction (DTI) prediction plays a crucial role in drug development, impacting areas such as virtual screening, drug repurposing, and the identification of potential drug side effects. Despite significant efforts dedicated to improving DTI prediction, existing methods still struggle with the challenges posed by the high sparsity of DTI datasets and the complexity of capturing heterogeneous information in biological networks. To address these challenges, we propose a unified framework for DTI prediction based on a semantics-guided meta-path walk. Specifically, we first pre-train drug and protein embeddings to capture their semantic information. This semantic information is then leveraged to guide a meta-path-based random walk on the biological heterogeneous network, generating sequences of interactions. These sequences are used to compute embedding features via a heterogeneous skip-gram model, which are subsequently fed into downstream tasks to predict DTIs. SGMDTI achieves substantial performance improvement over other state-of-the-art methods for drug–target interaction prediction. Moreover, it excels in the cold-start scenario, which is often a challenging case in DTI prediction. These results indicate the effectiveness of our approach in predicting drug-target interactions.Experimental datasets and experimental codes can be found in https://github.com/HYLPRC/SGMDTI

## 1 Introduction

The identification of drug–target interactions (DTI) plays a crucial role in various applications of drug development, such as lead discovery, drug repurposing, and the detection of off-target effects [1]. However, traditional biological experiments to detect DTI are often expensive and time-consuming [2]. Over the past few decades, numerous computational approaches have been developed to accelerate DTI prediction, helping guide in vivo validation and significantly advancing drug discovery and repositioning [3].

Computational methods for DTI prediction can be broadly classified into three categories: docking-based, ligand-based, and hybrid methods [4]. Structurebased virtual screening relies on molecular docking simulations, which become impractical if the three-dimensional structure of the target protein is unknown [5]. Ligand-based virtual screening, which predicts new active molecules based on known active ones, suffers when the number of known active molecules is insufficient [6]. Hybrid methods attempt to address these shortcomings and can be further divided into similarity-based, pharmacological feature-based, and network-based methods [7]. Similarity-based methods assume ‘guilt by association,’ while pharmacological feature-based methods focus on learning molecular features. Network-based methods, using binary and heterogeneous networks, aim to enhance DTI prediction by integrating various data types [8].

In complex biological systems, interactions between drugs and targets are influenced by numerous factors [9]. As a result, machine learning models based on a single data source often fail to capture the full complexity of molecular mechanisms underlying DTI, limiting predictive performance [10]. With advancements in biotechnology and the accumulation of biomedical data, integrating heterogeneous data sources is increasingly recognized as crucial for improving DTI predictions [11]. However, these approaches may still overlook the inherent diversity of biological networks. A common strategy for addressing this issue is the use of meta-paths, which represent sequences of node types connected by relationship types, effectively encapsulating biological principles within heterogeneous networks [12].

Despite the promise of meta-path-based approaches like SGCL-DTI [13] and GSL-DTI [14], these methods often fail to account for how the intrinsic properties of drugs and proteins influence the generated paths. To address these limitations, we propose SGMDTI, a semantic-guided meta-path method for predicting drug-target interactions. The flowchart of SGMDTI is as shown in Fig. 1. SGMDTI integrates semantic information-guided teleportation, incorporating drug and protein attributes into the meta-path generation process. The generated paths are used to create embedding vectors through a heterogeneous Skip-gram model, which are then employed with an XGBoost classifier to predict drug-protein associations. In multiple experimental settings, our method surpasses baseline approaches, including feature-based, end-to-end machine learning, and other network-based methods. We also explore the impact of various meta-path types on DTI prediction and investigate effective strategies for making more accurate inferences using SGMDTI. These results demonstrate SGMDTI’s robustness and scalability, establishing its efficacy in DTI prediction across a range of experimental setups.

**Fig. 1.**
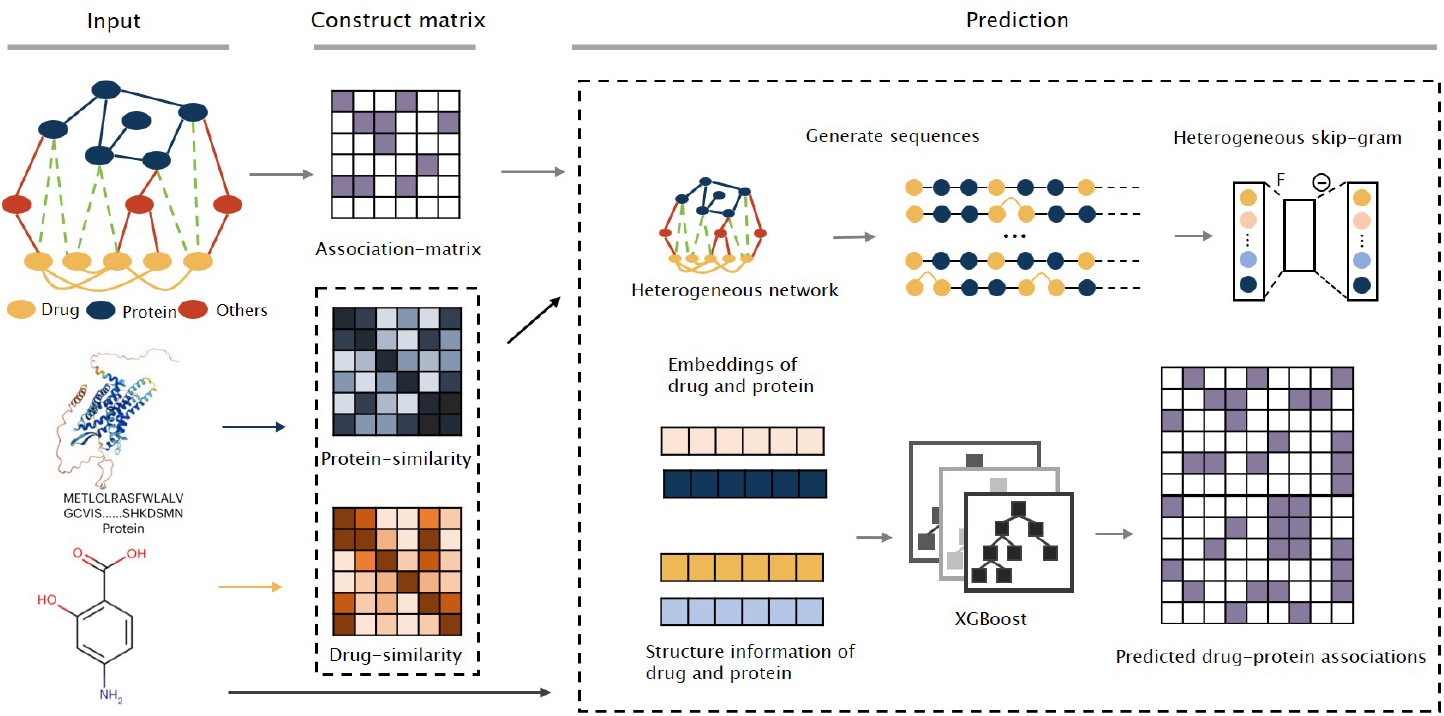
Flowchart of SGMDTI

## 2 Results

To rigorously evaluate the performance of the proposed model, SGMDTI is compared the performance of with the several baseline methods including DeepConVDTI [15], Moltrans [16], GNN-CPI [17], DeepEmbedding-DTI [18], PADMEECFP [19], metapath2vec [20], Multi_DTI [21], KGE_NFM [22], GIFDTI [23], Drugban [24], SGCL-DTI [13], GSL-DTI [14]. For detailed information on the baseline methods, please refer to the supplementary materials.

### 2.1 Performance evaluation of the proposed model

In this study, a 10-fold cross-validation approach was employed. Negative samples were constructed by randomly selecting drug-protein pairs with unknown associations. Specifically, the number of negative samples was ensured to match the number of positive interactions. Rather than using a fixed number of positive and negative samples in each fold, the entire dataset (both positive and negative samples) was randomly shuffled before performing cross-validation. In each fold, one subset was used as the test set, while the remaining nine subsets were used as the training set. This approach ensured a balanced evaluation and minimized potential bias during model training and testing. The result is shown in Fig. 2, with more details available in Supplementary Table S3.

**Fig. 2.**
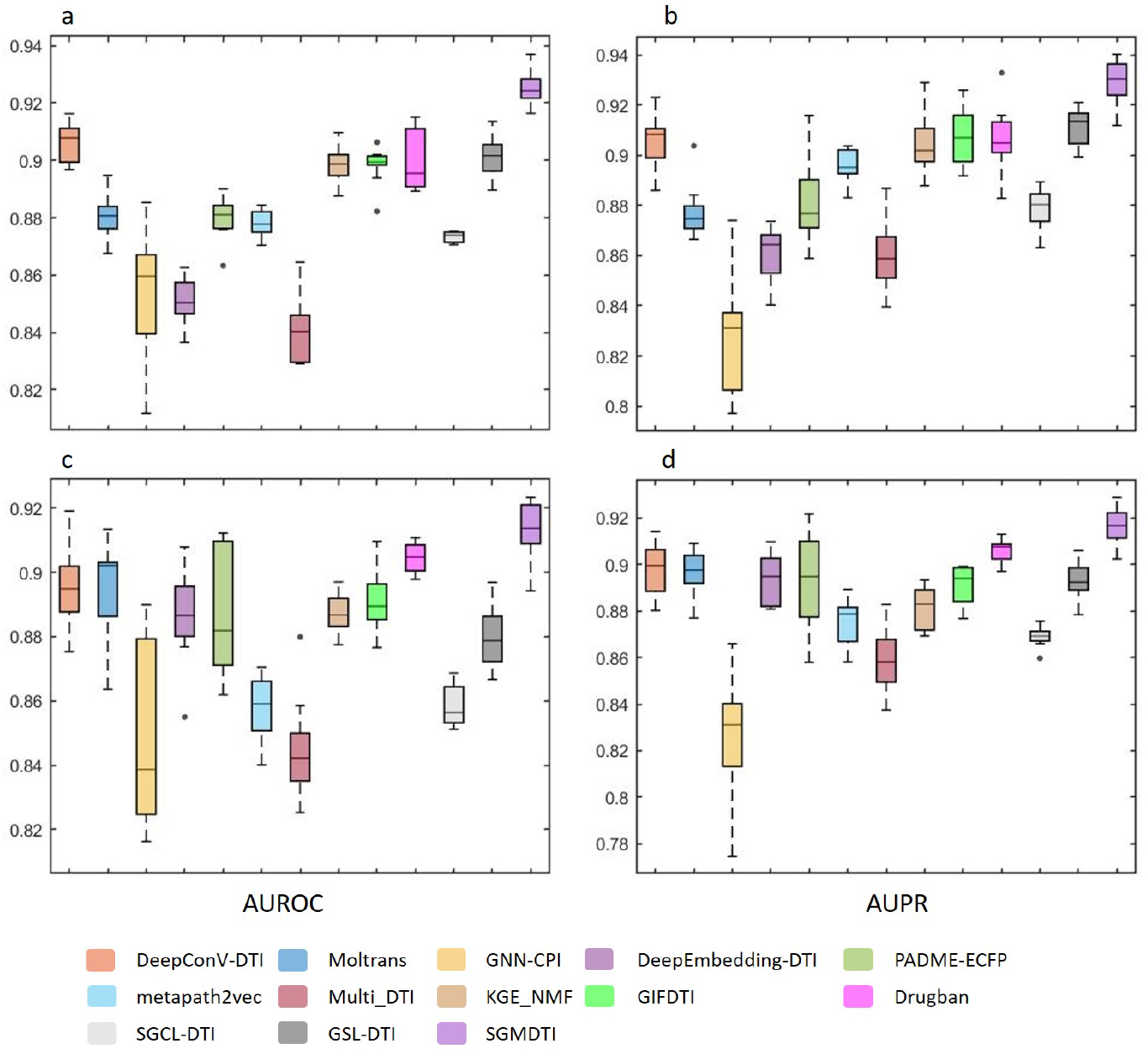
Results of 10-fold cross-validation. Subplots (a) and (b) show the AUC and AUPR for the Hetero-A dataset, respectively. Subplots (c) and (d) represent the AUC and AUPR for the Hetero-B dataset, respectively.

The results show that our proposed SGMDTI model outperforms all other models in both datasets, achieving the highest AUC and AUPR scores. On the Hetero-A dataset, SGMDTI achieved the AUC of 0.9250 and the AUPR of 0.9292, which are the highest among all models. When compared to the second-best model, DeepConv-DTI, SGMDTI shows an improvement of 0.0190 in AUC and 0.0227 in AUPR. On the Hetero-B dataset, SGMDTI similarly performed well, with an AUC of 0.9128 and an AUPR of 0.9168. Compared to the second-best model, Drugban, SGMDTI achieved an increase of 0.0085 in AUC and 0.0106 in AUPR.

These comparisons demonstrate that the SGMDTI model consistently achieves superior performance across different datasets. The improvements in AUC and AUPR highlight the effectiveness of integrating semantic information-guided meta-path generation in enhancing the predictive power for drug-target interaction.

### 2.2 Ablation study

To evaluate the contribution of each component within the SGMDTI framework, we conducted an ablation study by testing various model variants. This approach enabled us to isolate the effect of each component and gain a deeper understanding of its role in the overall performance of our method. The results of these experiments are presented in Fig. 3.

**Fig. 3.**
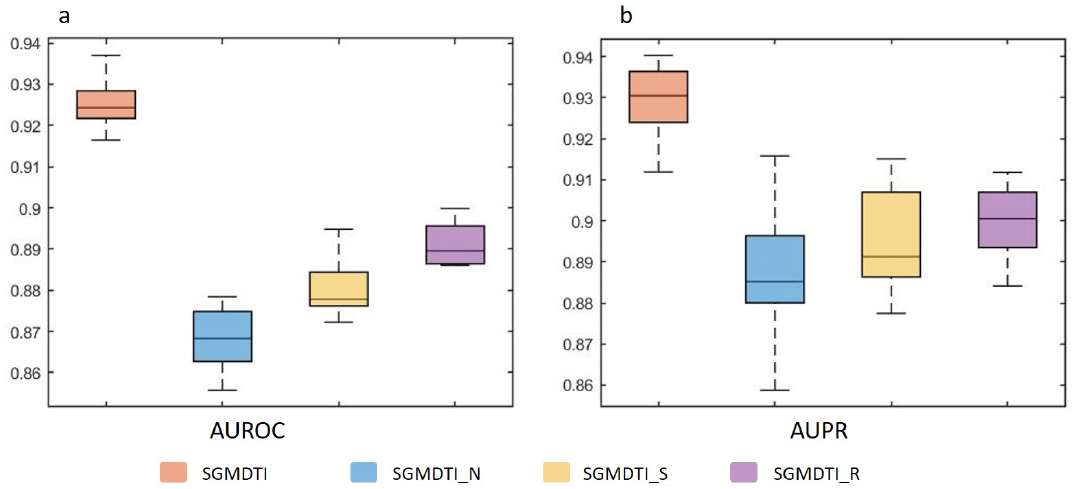
Ablation study results showing the impact of individual components on SGMDTI’s prediction performance

- **SGMDTI**: Our proposed model, which incorporates all components.
- **SGMDTI_N**: A variant where meta-path walking is omitted.
- **SGMDTI_S**: A variant that excludes the use of semantic information for guidance.
- **SGMDTI_R**: A variant where the heterogeneous Skip-gram model is not utilized.

### 2.3 Parametric analysis

To evaluate the influence of key parameters in the SGMDTI model, we performed the grid search experiment. The results are illustrated in Fig. 4. Two parameters were analyzed during this process:

**Fig. 4.**
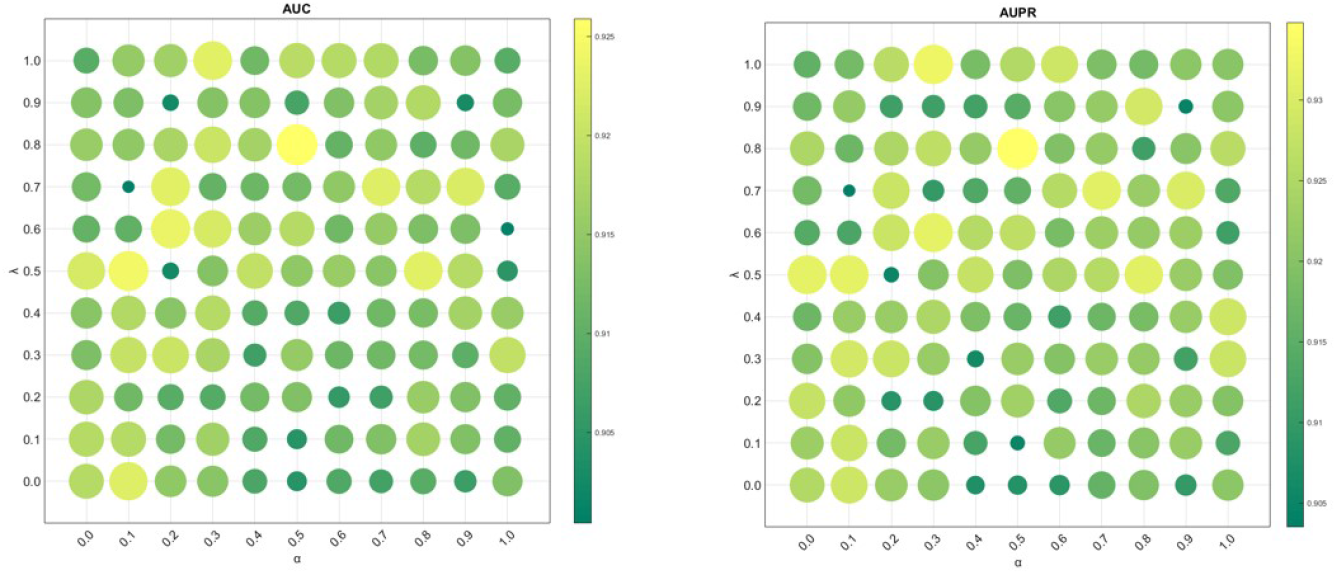
Parameter Sensitivity Analysis for *α* and *λ* in SGMDTI

(*α*): The threshold used for establishing edges in the attribute similarity network. (*λ*): The threshold used to balance between semantic-guided teleportation and meta-path walking.

The grid search explored parameter values within the range of [0, 1], with increments of 0.1. The results indicate that the SGMDTI model achieves optimal performance when *α* is set to 0.5 and *λ* is set to 0.8.

### 2.4 Cold start for drug and protein

In addition to the previous analyses, we conducted experiments under cold-start conditions for both drugs and proteins. The cold-start setting evaluates the model’s ability to predict interactions involving drugs and proteins that were not present in the training data. The results of these experiments, as shown in Fig. 5. For the cold start for drugs scenario, SGMDTI achieved the highest AUC of 0.8058 and AUPR of 0.8359. Compared to the second-best model, Muti_DTI (AUC: 0.8009, AUPR: 0.8126), SGMDTI shows an improvement of 0.0049 in AUC and 0.0233 in AUPR. The significant improvement in AUPR indicates that SGMDTI captures the true positive interactions more effectively than other models in the cold start drug scenario. For the cold start for proteins scenario, SGMDTI also achieves the highest performance with an AUC of 0.8181 and an AUPR of 0.8225. Compared to the second-best model, SGMDTI improves the AUC by 0.0077 and maintains a comparable AUPR. These results demonstrate that our proposed SGMDTI model consistently outperforms other models in both cold start scenarios.

**Fig. 5.**
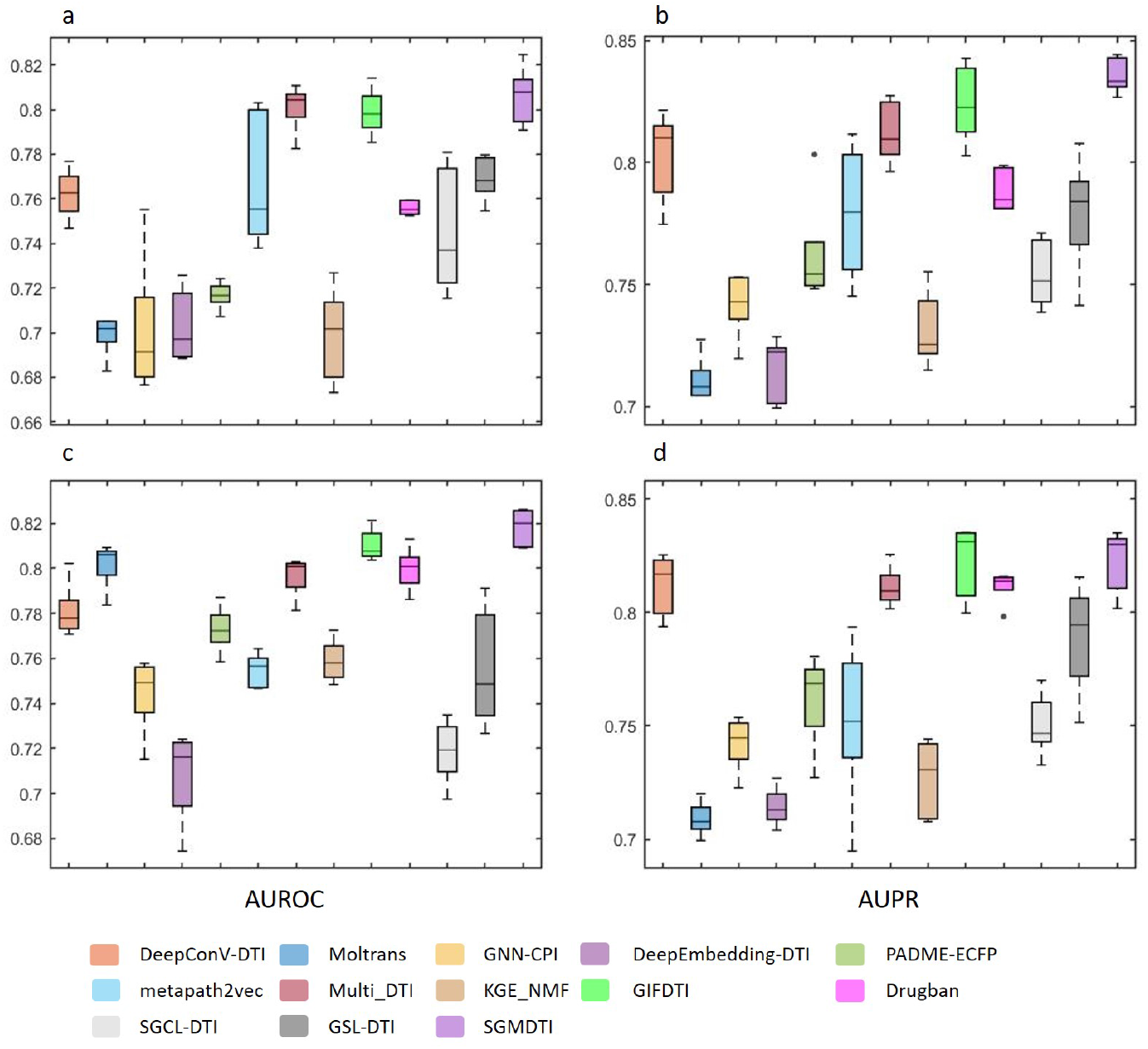
Comparison of Drug Cold-Start Performance Between SGMDTI and Baseline Methods

### 2.5 Meta-path-based experiment results

To further evaluate the performance of SGMDTI, we conducted an experiment by selectively removing certain meta-paths from the model to assess their impact on predictive accuracy. The result is as shown in Table 1.

**Table 1.** Comparison of different model variants based on metapath.

| Model | AUROC | AUPRC |
| --- | --- | --- |
| Without drug-drug | 0.9081 | 0.9106 |
| Without drug-se-drug | 0.9113 | 0.9213 |
| Without drug-disease-drug | 0.8973 | 0.9031 |
| Without protein-protein | 0.9112 | 0.9168 |
| Without protein-disease-protein | 0.8852 | 0.8968 |
| **Proposed model** | **0.9220** | **0.9291** |

The results show that removing the protein-disease-protein meta-path had the greatest impact, with the AUC dropping from 0.9220 to 0.8852. This indicates that this meta-path plays a crucial role in capturing interactions between proteins and diseases, significantly contributing to the model’s overall performance. In contrast, removing the protein-protein and drug-se-drug meta-paths had minimal impact on performance. When the protein-protein relations were removed, the AUC only dropped by 0.0108, while removing the drug-se-drug relations resulted in a similar AUC decrease of 0.0107. The minimal impact of these two meta-paths on the model’s performance could be due to their abundance in the network, which introduces a large amount of data. However, this also likely introduces some noise that is unrelated to the entity research focus, further diluting their overall contribution to the model’s predictive ability. In addition, removing any of these meta-paths results in a decline in predictive accuracy, confirming the importance of integrating diverse relationships in the model.

### 2.6 Casestudy

We conducted two case studies to further validate the performance of SGMDTI in predicting drug-target interactions (DTI) related to COVID-19. Specifically, we focused on two proteins that have been linked to COVID-19: Apolipoprotein E (P02649) [25] and Angiotensin-converting enzyme 2 (Q9BYF1) [26]. For each protein, we performed a Top 5 prediction of potential drug candidates. These predictions offer insights into possible drug repositioning opportunities for combating COVID-19. Detailed results of these case studies are provided in the supplementary materials.

## 3 Methods

### 3.1 Benchmark datasets

In this study, two benchmark datasets were used to evaluate the performance of our model as well as state-of-the-art models for drug-target interaction (DTI) prediction. The first dataset, denoted as Hetero-A, is derived primarily from DrugBank and contains DTI data as well as additional information, including drugs, proteins, diseases, and drug side effects [27]. The second dataset, denoted as Hetero-B, is compiled from BioSnap along with related disease datasets [28]. Detailed descriptions and statistics of these datasets can be found in the supplementary materials, and the data can be downloaded from https://github.com/HYLPRC/SGMDTI/SGMDTI-master/data.

### 3.2 Drug and protein semantic similarities

In this study, we integrate semantic information into biological heterogeneous networks to achieve drug-target interaction prediction through semantically guided meta-path embeddings. The semantic information of drugs and proteins primarily consists of molecular structure characteristics of drugs and amino acid sequence information of proteins. Based on the features of drug and protein ontologies, we calculate the semantic similarity between drug-drug and proteinprotein pairs. The detailed process of calculating these similarities is described as follows.

#### Drug feature

Each drug’s SMILES string *S* is first converted into a molecular graph *G* = (*V, E*), where *V* represents atoms and *E* represents chemical bonds [29], using RDKit:

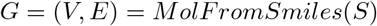

For each graph *G*, nodes (atoms) and edges (bonds) are assigned features. Node features 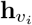 and edge features 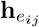 are extracted using predefined functions:

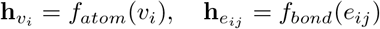

The GIN (Graph Isomorphism Network) model is then applied to generate graph embeddings by aggregating neighborhood information across multiple layers. The message-passing mechanism of GIN is as follows:

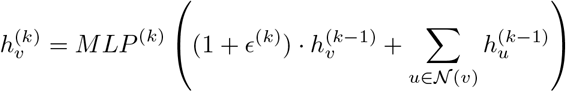

where 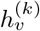 is the embedding of node *v* after the *k*-th layer, 𝒩 (*v*) represents the neighbors of node *v, ϵ*^(*k*)^ is a learnable parameter that controls the relative weight between the node’s own features and those of its neighbors, and *MLP* ^(*k*)^ is a multi-layer perceptron for non-linear transformation.

After the graph convolutions, a Readout operation pools the node embeddings into a single molecular representation:

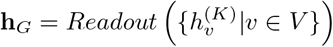

We use the pretrained gin_supervised_contextpred model to extract highquality molecular embeddings, leveraging context prediction tasks. The molecular embedding **h**_*G*_ is then obtained from the pretrained GIN model:

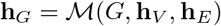

#### Protein feature

In this study, CTD (Composition, Transition, Distribution) descriptors were utilized to extract protein features [30]. These descriptors, which generate numerical features by calculating the composition, transition frequency, and distribution patterns of amino acids within the protein sequence, were employed to capture the global physicochemical properties of the proteins. By focusing on these properties, a comprehensive representation of the protein’s structural and functional characteristics was obtained. This approach enabled the incorporation of detailed sequence-level information into the predictive model, contributing to a more robust and accurate characterization of the proteins.

#### Semantic similarities

In this study, the internal similarity between drugs and proteins was calculated separately [8]. For drugs, the Euclidean distance between each pair of drugs was first computed using their feature vectors. Given two drugs with feature vectors 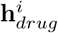 and 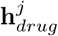, the Euclidean distance was defined as:

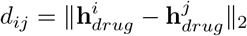

The distances were then converted into similarity scores using the formula:

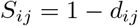

where *S*_*ij*_ represents the similarity between drug *i* and drug *j*, and *d*_*ij*_ is their Euclidean distance. A similar process was applied to calculate the similarity between proteins.

### 3.3 Semantic-guided meta-path

In this study, a semantic-guided meta-path method was employed to generate node sequences in the heterogeneous network. During the walk process, when the current walk was at node *v*_*i*_, the next step could transition to any neighboring node *v*_*j*_ ∈ *N* (*v*_*i*_). The probability of choosing *v*_*j*_ as the next node *P* (*v*_*i*+1_ = *v*_*j*_ | *v*_*i*_) was computed using the softmax function:

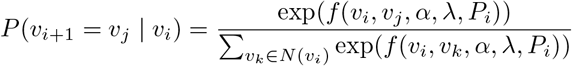

In this formula, *f* (*v*_*i*_, *v*_*k*_, *α, λ, P*_*i*_) is a feature function that measures the relationship between the current node *v*_*i*_ and the neighboring node *v*_*j*_ along the meta-path *P*_*i*_. The term *α* is a parameter for establishing edges in the attribute similarity network.The term *λ* is a parameter from the semantic network and meta-path that adjusts the transition weight from node *v*_*i*_ to *v*_*j*_. The set *N* (*v*_*i*_) represents all neighboring nodes of the current node *v*_*i*_.

Once the sequences are generated, Heterogeneous Skip-gram (HSG) model is employed to extract the feature of drug and protein [31]. To improve performance, HSG adopted a negative sampling strategy that took the heterogeneity of nodes into consideration. Specifically, during negative sampling, the model ensured that negative samples were drawn from the same type of nodes as the current node.Detailed information about SGMDTI can be found in the supplementary materials.

In this study, XGBoost was utilized to perform classification tasks based on the embeddings extracted from the semantic-guided meta-path model [32]. The feature vectors representing drug-target pairs were used as input to the XGBoost classifier, and the model was trained to predict the likelihood of interaction between drugs and targets. Through XGBoost’s iterative boosting process, the model effectively learned to distinguish between interacting and non-interacting pairs, providing accurate predictions for drug-target interaction tasks.

## 4 Conclusion

In this study, a novel approach for drug-target interaction prediction was proposed by integrating semantic information into a heterogeneous biological network. Through the use of semantically guided meta-paths, meaningful sequences were generated to capture both the structural and semantic relationships within the network. A Heterogeneous Skip-gram (HSG) model was employed, which efficiently extracted node embeddings while considering the heterogeneity of nodes such as drugs and proteins. By leveraging negative sampling specific to node types, the quality of the embeddings was enhanced, making them suitable for downstream tasks like drug-target interaction prediction. The experimental results demonstrated the effectiveness of this approach, outperforming baseline models in several metrics. This method not only improved predictive accuracy but also provided insights into the complex relationships between biological entities, paving the way for more advanced applications in drug discovery and repositioning.

## Acknowledgments

This work was carried out in part using computing resources at the High Performance Computing Center of Central South University.

## References

1. Ying Luo, Xiaotong Zhao, Mu Zhou, Yong Fang, Yong Li, Xu Zhang, Yu Peng, Xian He, and Hao Deng. A network integration approach for drug-target interaction prediction and computational drug repositioning from heterogeneous information. Nature Communications, 8:1–13, 2017.

2. Yoshihiro Yamanishi, Mitsuo Araki, Alex Gutteridge, Wataru Honda, and Minoru Kanehisa. Prediction of drug-target interaction networks from the integration of chemical and genomic spaces. Bioinformatics, 24(issue supplement 1):i232–i240, 2008.

3. Hakime Öztürk, Elif O. Olmez, and Arzucan Özgür. Deepdta: deep drug-target binding affinity prediction. Bioinformatics, 34(issue supplement 13):i821–i829, 2018.

4. Sungah Lim et al. A review on compound-protein interaction prediction methods: data, format, representation and model. Computational and Structural Biotechnology Journal, 19:1541–1556, 2021.

5. Jan Sieg, Fabian Flachsenberg, and Matthias Rarey. In need of bias control: evaluating chemical data for machine learning in structure-based virtual screening. Journal of Chemical Information and Modeling, 59:947–961, 2019.

6. Kai-Yue Gao et al. Interpretable drug target prediction using deep neural representation. In Proceedings of the International Joint Conference on Artificial Intelligence (IJCAI), pages 3371–3377, 2018.

7. Markus Bredel and Edgar Jacoby. Chemogenomics: an emerging strategy for rapid target and drug discovery. Nature Reviews Genetics, 5:262–275, 2004.

8. Dongmin Bang, Sangsoo Lim, Sangseon Lee, and Sun Kim. Biomedical knowledge graph learning for drug repurposing by extending guilt-by-association to multiple layers. Nature Communications, 14(1):3570, 2023.

9. Yanyi Chu, Aman Chandra Kaushik, Xiangeng Wang, Wei Wang, Yufang Zhang, Xiaoqi Shan, Dennis Russell Salahub, Yi Xiong, and Dong-Qing Wei. Dti-cdf: a cascade deep forest model towards the prediction of drug-target interactions based on hybrid features. Briefings in Bioinformatics, 22(1):451–462, 2021.

10. Ran Zhang, Zhanjie Wang, Xuezhi Wang, Zhen Meng, and Wenjuan Cui. Mhtandti: Metapath-based hierarchical transformer and attention network for drug– target interaction prediction. Briefings in Bioinformatics, 24(2):bbad079, 2023.

11. Brian Walsh, Sameh K Mohamed, and Vít Nováček. Biokg: A knowledge graph for relational learning on biological data. In Proceedings of the 29th ACM International Conference on Information & Knowledge Management, pages 3173–3180, 2020.

12. Hongzhun Wang, Feng Huang, Zhankun Xiong, and Wen Zhang. A heterogeneous network-based method with attentive meta-path extraction for predicting drug– target interactions. Briefings in Bioinformatics, 23(4):bbac184, 2022.

13. Yilong Li, Guojun Qiao, Xianzhe Gao, and Guangyu Wang. Supervised graph co-contrastive learning for drug–target interaction prediction. Bioinformatics, 38(10):2847–2854, 2022.

14. Zixuan E, Guojun Qiao, Guangyu Wang, and Yilong Li. Gsl-dti: Graph structure learning network for drug-target interaction prediction. bioRxiv, 2024.

15. Inhye Lee, Jinwook Keum, and Hojung JP Nam. Deepconv-dti: Prediction of drug-target interactions via deep learning with convolution on protein sequences. PLOS Computational Biology, 15(6):e1007129, 2019.

16. Kun Huang, Cao Xiao, Lucas M Glass, and Jimeng Sun. Moltrans: molecular interaction transformer for drug–target interaction prediction. Bioinformatics, 37(6):830–836, 2021.

17. Masashi Tsubaki, Kentaro Tomii, and Jun Sese. Compound–protein interaction prediction with end-to-end learning of neural networks for graphs and sequences. Bioinformatics, 35(2):309–318, 2019.

18. Wenlin Chen, Gong Chen, Lei Zhao, and Chuan-Yu Chen. Predicting drug–target interactions with deep-embedding learning of graphs and sequences. The Journal of Physical Chemistry A, 125(25):5633–5642, 2021.

19. Qianqian Feng, Evgeniya Dueva, Artem Cherkasov, and Martin Ester. Padme: A deep learning-based framework for drug-target interaction prediction. arXiv preprint 1807.09741, 2018.

20. Yuxiao Dong, Nitesh V Chawla, and Ananthram Swami. metapath2vec: Scalable representation learning for heterogeneous networks. In Proceedings of the 23rd ACM SIGKDD international conference on knowledge discovery and data mining, pages 135–144, 2017.

21. Dingxian Zhou, Zenghui Xu, Wei Li, Xiaohong Xie, and Shuguang Peng. Multidti: drug–target interaction prediction based on multi-modal representation learning to bridge the gap between new chemical entities and known heterogeneous network. Bioinformatics, 37(23):4485–4492, 2021.

22. Qianqian Ye et al. A unified drug–target interaction prediction framework based on knowledge graph and recommendation system. Scientific Reports, 12(1):6775, 2021.

23. Qian Zhao et al. Gifdti: Prediction of drug-target interactions based on global molecular and intermolecular interaction representation learning. bioRxiv, 2022.

24. Ping Bai, Filip Miljković, Brian John, and Haifeng Lu. Interpretable bilinear attention network with domain adaptation improves drug–target prediction. Nature Machine Intelligence, 5(2):126–136, 2023.

25. Feng Chen, Qiongwei Ke, Wenyan Wei, Lili Cui, and Yan Wang. Apolipoprotein e and viral infection: Risks and mechanisms. Molecular Therapy-Nucleic Acids, 2023.

26. Markus Hoffmann, Hannah Kleine-Weber, Simon Schroeder, Nadine Krüger, Tanja Herrler, Sandra Erichsen, Tobias S. Schiergens, Georg Herrler, Nai-Huei Wu, Andreas Nitsche, Marcel A. Müller, Christian Drosten, and Stefan Pöhlmann. Sarscov-2 cell entry depends on ace2 and tmprss2 and is blocked by a clinically proven protease inhibitor. Cell, 181(2):271–280.e8, 2020.

27. David S Wishart, Craig Knox, An Chi Guo, Dean Cheng, Savita Shrivastava, Dan Tzur, Bijaya Gautam, and Murtaza Hassanali. Drugbank: a knowledgebase for drugs, drug actions and drug targets. Nucleic acids research, 36(Suppl_1):D901– D906, 2008.

28. Sukannya Purkayastha, Ishani Mondal, Sudeshna Sarkar, Pawan Goyal, and Jitesh K Pillai. Drug-drug interactions prediction based on drug embedding and graph auto-encoder. In 2019 IEEE 19th International Conference on Bioinformatics and Bioengineering (BIBE), pages 547–552. IEEE, 2019.

29. Weihua Hu, Bowen Liu, Joseph Gomes, Marinka Zitnik, Percy Liang, Vijay Pande, and Jure Leskovec. Strategies for pre-training graph neural networks. In International Conference on Learning Representations, 2020.

30. Anton Meinhart, Tomislav Kamenski, Sabine Hoeppner, Sonja Baumli, and Patrick Cramer. A structural perspective of ctd function. Genes & development, 19(12):1401–1415, 2005.

31. Peng Wu, Conghui Zheng, and Li Pan. A unified generative adversarial learning framework for improvement of skip-gram network representation learning methods. IEEE Transactions on Knowledge and Data Engineering, 35(1):45–58, 2021.

32. Tianqi Chen and Carlos Guestrin. Xgboost: A scalable tree boosting system. In Proceedings of the 22nd acm sigkdd international conference on knowledge discovery and data mining, pages 785–794, 2016.

